## Supplementary information for "Distributions of extinction times from fossil ages and tree topologies: the example of mid-Permian synapsid extinctions"

#### SI-1 Previous approaches

##### SI-1.1 Branch-by-branch approaches

We present here several previous approaches which deal with each taxon independently of the rest of the tree, in the sense that in order to estimate the extinction time of a given (extinct) taxon, they only require the number $n$ of its fossils and their ages $f_1$, $f_2$, ..., $f_n$. We assume that the fossil ages are given in chronological order, so $f_1$ and $f_n$ are the most ancient and most recent fossil ages respectively.

The approach of Strauss and Sadler (1989) makes the assumption that the fossil ages of a given extinct taxon are uniformly distributed into a time interval of unknown bounds, which are its speciation and extinction ages. By using properties of uniformly distributed samples, they provided, amongst others, an explicit expression of confidence upper bound of the extinction age of a given taxon at order $o$, namely $f_n + (1-o)^{-\frac{1}{n-1}}(f_n - f_1)$.

McInerny *et al.* (2006) proposed a method to infer extinction times from sighting records under a uniformity assumption close to that Strauss and Sadler (1989). Let us briefly expose their approach by transposing it into the geological case. The method assumes that the probability of fossilization per unit of time is constant and estimates it as $\frac{n-1}{f_n-f_1}$ (in the case where the first fossil age is used as start time, the number of fossiliferous horizons considered reduce to $n-1$ and let us assume here that at most a single fossiliferous horizon is observed per unit of time). The probability for the extinction time to be posterior to any time $T > f_n$ is that of not observing any fossil between $f_n$ and time $T$ given that the considered lineage does not go extinct before $T$, which follows a geometric distribution, $\left(1 - \frac{n-1}{f_n-f_1}\right)^{T-f_n}$, and from which we can get a confidence upper bound at a given order $o$: $f_n + \frac{\log(1-o)}{\log\left(1-\frac{n-1}{f_n-f_1}\right)}$. A continuous version of this approach is basically obtained by replacing the geometric distribution by an exponential distribution in the formula just above. The probability for the the extinction time to be posterior to a time $T$ becomes $\exp\left(-\frac{n-1}{f_n-f_1}(T-f_n)\right)$ and the confidence upper bound at order $o$ is then $f_n - \frac{\log(1-o)}{\left(\frac{n-1}{f_n-f_1}\right)}$. Note that $\frac{n-1}{f_n-f_1}$ is now interpreted as the rate of a Poisson process modelling the fossilization (and we no longer have to assume that at most a single fossiliferous horizon is observed per unit of time).

Bradshaw *et al.* (2012) proposed a method based on that of McInerny *et al.* (2006) in order to estimate extinction times from the fossil record. They first consider a weighted combination of the results provided by the method of McInerny *et al.* (2006) on the $k$ most recent fossils for $k = 1$ to $n-1$ with weights inversely proportional to $f_n - f_{n-k}$, in order to give more weight to the most recent fossils. They also take into account the uncertainty in dating fossils by resampling the fossil ages from Gaussian distributions and by considering 95% confidence limits.

Alroy (2014) proposed a Bayesian iterative approach to infer extinction date – see Solow (2016) and Alroy (2016) for a discussion of this method. Briefly, the time after the most recent fossil age is divided into equal subintervals indexed on $1, 2, \ldots$. Under the notations of Alroy (2014), let $p_s$ be the probability that a particular interval contains a fossil age given that the taxon is not extinct at this time interval, $E$ be a prior probability of extinction and $\epsilon_t$ be the posterior probability of extinction before the $t^{\text{th}}$ interval (Alroy, 2014, proposed different ways to estimate the two first quantities). The method iteratively computes the posterior probability of extinction before all intervals $t$ by setting $\epsilon_1 = \frac{E}{E+(1-E)(1-p_s)}$ and, for all $t > 1$, $\epsilon_t = \frac{\epsilon_{t-1}+(1-\epsilon_{t-1})E}{\epsilon_{t-1}+(1-\epsilon_{t-1})E+(1-\epsilon_{t-1})(1-E)(1-p_s)}$. The confidence upper bound of the extinction date at order $o$ is the upper bound of the last interval $t$ such that $\epsilon_t \leq o$.

##### SI-1.2 A basic global method

The approach of McInerny *et al.* (2006) and its continuous version can be adapted in order to take into account the whole tree by considering all the fossiliferous horizons for the estimation of the probability or the rate of fossilization. This global estimation is performed by considering all the differences between the ages of the successive fossils of the phylogenetic tree. In the continuous case, these time differences are assumed to be exponentially distributed with the rate of the Poisson process modelling the fossilization, and we consider its maximum likelihood estimate $\hat{\varphi}$. The 'global' fossilization rate estimate $\hat{\varphi}$ is next used to get the confidence upper bound at order $o$ of the extinction time of any extinct taxa of the tree as above, i.e., $f_n - \frac{\log(1-o)}{\hat{\varphi}}$ in the continuous case.

#### SI-2 Probability that a subset of extinct taxa goes extinct before another

Let $\mathcal{T}$ be a tree with fossils (without its divergence times), $\mathbf{f}$ be a fossil age vector and $A$ and $B$ be two disjoint subsets of extinct taxa of $\mathcal{T}$ (e.g., two mutually exclusive extinct clades of $\mathcal{T}$). We aim here to compute the probability density $\mathbf{P}(\mathcal{T}, \mathbf{f}, \eta_A \leq \eta_B)$, which is the joint probability density of the data $(\mathcal{T}, \mathbf{f})$ and that the taxa of $A$ go extinct before those of $B$.

We shall consider the situation where the data are provided as a set $\Omega$ of equiparsimonious trees and that the fossil ages are just bounded by geological intervals summarized in the product-set $\mathbf{I}$. The joint probability density above conditioned on the whole dataset is

$$\frac{\sum_{\mathcal{T}\in\Omega}\int_{\mathbf{I}}\mathbf{P}(\mathcal{T}, \mathbf{f}, \eta_A \leq \eta_B)\mathrm{d}\mathbf{f}}{\sum_{\mathcal{T}\in\Omega}\int_{\mathbf{I}}\mathbf{P}(\mathcal{T}, \mathbf{f})\mathrm{d}\mathbf{f}} = \frac{\int_{\mathbf{I}}\sum_{\mathcal{T}\in\Omega}\mathbf{P}(\mathcal{T}, \mathbf{f}, \eta_A \leq \eta_B)\mathrm{d}\mathbf{f}}{\int_{\mathbf{I}}\sum_{\mathcal{T}\in\Omega}\mathbf{P}(\mathcal{T}, \mathbf{f})\mathrm{d}\mathbf{f}}. \tag{1}$$

Let us first show how to compute $\mathbf{P}(\mathcal{T}, \mathbf{f}, \eta_A \leq \eta_B)$. We recall that for all extinct taxa $n$, $\ell_{\mathbf{f},n}$ stands for the most recent fossil age of $n$ and we put $\ell_{\mathbf{f},A\cup B}$ for the most recent age of fossils of $A \cup B$, i.e., $\ell_{\mathbf{f},A\cup B} = \max_{n\in A\cup B}\ell_{\mathbf{f},n}$. The probability density $\mathbf{P}(\mathcal{T}, \mathbf{f}, \eta_A \leq \eta_B)$ can be obtained by integrating over all the possible ages $t$ of extinction of $B$, the joint probability density that $B$ goes extinct exactly at $t$ and $A$ goes extinct before $t$. Note that the possible ages $t$ such that $B$ goes extinct exactly at $t$ and $A$ goes extinct before $t$ run from $\ell_{\mathbf{f},A\cup B}$ to the present time $T$. Moreover, the probability density that $B$ goes extinct exactly at $t$ is the sum over all the taxa $m$ of $B$ of the probability densities that $m$ goes extinct exactly at $t$ and all the other taxa of $B$ go extinct before $t$. It follows that the probability density $\mathbf{P}(\mathcal{T}, \mathbf{f}, \eta_A \leq \eta_B)$ can be written

$$\begin{aligned}
\mathbf{P}(\mathcal{T}, \mathbf{f}, \eta_A \leq \eta_B) &= \int_{\ell_{\mathbf{f},A\cup B}}^{T} \mathbf{P}(\mathcal{T}, \mathbf{f}, \eta_A \leq t, \eta_B = t)\mathrm{d}t \\
&= \int_{\ell_{\mathbf{f},A\cup B}}^{T} \sum_{m\in B} \mathbf{P}(\mathcal{T}, \mathbf{f}, \eta_A \leq t, \eta_{B\setminus\{j\}} \leq t, \eta_m = t)\mathrm{d}t \\
&= \frac{\mathbf{P}(\mathcal{T}, \mathbf{f})\sum_{m\in B}\int_{\ell_{\mathbf{f},A\cup B}}^{T}\left(\prod_{n\in A\cup B\setminus\{m\}}\mathbf{P}(0, t-\ell_{\mathbf{f},n})\right)\mathbf{D}(t-\ell_{\mathbf{f},m})\mathrm{d}t}{\prod_{n\in A\cup B}\mathbf{P}(0, T-\ell_{\mathbf{f},n})} \\
&= \frac{\mathbf{P}(\mathcal{T}, \mathbf{f})\mathcal{I}_{\mathbf{f},A,B}}{\prod_{n\in A\cup B}\mathbf{P}(0, T-\ell_{\mathbf{f},n})},
\end{aligned}$$

where $\mathcal{I}_{\mathbf{f},A,B} = \sum_{m\in B}\int_{\ell_{\mathbf{f},A\cup B}}^{T}\left(\prod_{n\in A\cup B\setminus\{m\}}\mathbf{P}(0, t-\ell_{\mathbf{f},n})\right)\mathbf{D}(t-\ell_{\mathbf{f},m})\mathrm{d}t$.

Since we assume here that the extant sampling parameter $\rho = 1$, Equations 1 of the main document give us that

$$\mathcal{I}_{\mathbf{f},A,B} = \sum_{m\in B} \int_{\ell_{\mathbf{f},A\cup B}}^{T} \left( \prod_{n\in A\cup B\setminus\{m\}} \frac{\alpha\beta(1-\mathrm{e}^{\omega(t-\ell_{\mathbf{f},n})})}{\beta-\alpha\mathrm{e}^{\omega(t-\ell_{\mathbf{f},n})}} \right) \frac{\mu(\beta-\alpha)^2\mathrm{e}^{\omega(t-t_m)}}{(\beta-\alpha\mathrm{e}^{\omega(t-t_m)})^2}\mathrm{d}t.$$

A tedious calculation leads to

$$\begin{aligned}\mathcal{I}_{\mathbf{f},A,B} = \beta(\beta-\alpha)\sum_{m\in B}\left(\sum_{n\in A\cup B\setminus\{m\}} \frac{(\beta-\alpha)\Upsilon_{\mathbf{f},m,n}}{1-\mathrm{e}^{\omega(\ell_{\mathbf{f},m}-\ell_{\mathbf{f},n})}} - \beta^{|A\cup B|-1}\right)\left[\frac{1}{\beta-\alpha\mathrm{e}^{\omega(t-\ell_{\mathbf{f},m})}}\right]_{\ell_{\mathbf{f},A\cup B}}^{T} \\ -(\beta-\alpha)^2\sum_{m\in B}\sum_{n\in A\cup B\setminus\{m\}} \frac{\Upsilon_{\mathbf{f},m,n}\mathrm{e}^{\omega(\ell_{\mathbf{f},m}-\ell_{\mathbf{f},n})}}{(1-\mathrm{e}^{\omega(\ell_{\mathbf{f},m}-\ell_{\mathbf{f},n})})^2}\left[\log\left(1+\frac{\alpha(1-\mathrm{e}^{\omega(\ell_{\mathbf{f},m}-\ell_{\mathbf{f},n})})}{\beta\mathrm{e}^{\omega(\ell_{\mathbf{f},m}-t)}-\alpha}\right)\right]_{\ell_{\mathbf{f},A\cup B}}^{T},\end{aligned} \quad (2)$$

where

$$\Upsilon_{\mathbf{f},m,n} = \prod_{k\in(A\cup B)\setminus\{m,n\}} \frac{\beta-\alpha\mathrm{e}^{\omega(\ell_{\mathbf{f},k}-\ell_{\mathbf{f},n})}}{1-\mathrm{e}^{\omega(\ell_{\mathbf{f},k}-\ell_{\mathbf{f},n})}}.$$

We thus have an explicit formula to compute $\mathbf{P}(\mathcal{T},\mathbf{f},\eta_A \leq \eta_B)$ which can be used in the importance sampling procedure presented in Appendix SI-3 to estimate the probability that the subset $A$ goes extinct before $B$. Namely, if $\mathbf{f}'_1, \ldots, \mathbf{f}'_N$ are $N$ iid samples of density $\mathrm{q}^\star$, the ratio of Equation 1 is estimated by

$$\begin{aligned}\frac{\frac{1}{N}\sum_{i=1}^{N}\frac{\sum_{\mathcal{T}\in\Omega}\mathbf{P}(\mathcal{T},\mathbf{f}'_i,\eta_A\leq\eta_B)}{\mathrm{q}^\star(\mathbf{f}'_i)}}{\int_{\mathbf{I}}\sum_{\mathcal{T}\in\Omega}\mathbf{P}(\mathcal{T},\mathbf{f})\mathrm{d}\mathbf{f}} &= \frac{1}{N}\sum_{i=1}^{N}\frac{\sum_{\mathcal{T}\in\Omega}\mathbf{P}(\mathcal{T},\mathbf{f}'_i,\eta_A\leq\eta_B)}{\sum_{\mathcal{T}\in\Omega}\mathbf{P}(\mathcal{T},\mathbf{f}'_i)} \\ &= \frac{1}{N}\sum_{i=1}^{N}\frac{\mathcal{I}_{\mathbf{f}'_i,A,B}}{\prod_{n\in A\cup B}\mathbf{P}(0,T-\ell_{\mathbf{f}'_i,n})}.\end{aligned}$$

### SI-3 A Metropolis-Hastings importance sampling procedure for the distribution of the extinction times of an arbitrary subset of taxa

We aim to compute the joint probability of a tree $\mathcal{T}$ with fossils (without its divergence times), a fossil age vector $\mathbf{f}$ and its origin time, and that a particular subset $S$ of (extinct at the present time) taxa of $\mathcal{T}$ goes extinct before a time $t$. Let us put $\eta_S$ for the extinction time of the taxa in $S$ and for all taxon $n$ of $S$, $\ell_{\mathbf{f},n}$ for the age of its most recent fossil with regard to the fossil age vector $\mathbf{f}$. The probability density of interest, $\mathbf{P}(\mathcal{T},\mathbf{f},\eta_n \leq t)$, reads

$$\mathbf{P}(\mathcal{T},\mathbf{f},\eta_S \leq t) = \mathbf{P}(\mathcal{T},\mathbf{f})\prod_{n\in S}\frac{\mathbf{P}(0,t-\ell_{\mathbf{f},n})}{\mathbf{P}(0,T-\ell_{\mathbf{f},n})}.$$

The ratio $\frac{\mathbf{P}(0,t-\ell_{\mathbf{f},n})}{\mathbf{P}(0,T-\ell_{\mathbf{f},n})}$ means that we replace the factor contribution of the unobservable tree ending the evolution of taxon $n$ in $\mathbf{P}(\mathcal{T},\mathbf{f})$, which is the probability that it goes extinct before the present time, i.e., $\mathbf{P}(0,T-\ell_{\mathbf{f},n})$, by $\mathbf{P}(0,t-\ell_{\mathbf{f},n})$, which is the probability that it goes extinct before $t$.

The computation of the probability density above holds in the case where both the topology and the fossil ages are exactly known. In practice, there is a lot of uncertainty over these two features. In particular, our dataset contains a set $\Omega$ of equiparsimonious topologies and the only information about the age of each fossil $f$ is provided as a geological age interval $\mathrm{I}_f$. Given that with a tree $\mathcal{T}\in\Omega$ and a vector of fossil ages $\mathbf{f}\in\mathbf{I}$, we are able to compute the probability densities $\mathbf{P}(\mathcal{T},\mathbf{f},\eta_S \leq t)$ for all times $t$, determining distributions from our dataset may be performed by summing over all the topologies and integrating over all the time intervals.

We assume prior distributions in which (i) all the topologies in $\Omega$ are equiprobable and (ii) the fossil ages are uniformly distributed over their time intervals. We put $\mathbf{I}$ for the product-set of the time intervals $\mathbf{I} = \bigotimes_f \mathrm{I}_f$. The joint probability density above conditioned on the whole dataset is

$$\frac{\sum_{\mathcal{T} \in \Omega} \int_{\mathbf{I}} \mathbf{P}(\mathcal{T}, \mathbf{f}, \eta_S \leq t)\mathrm{d}\mathbf{f}}{\sum_{\mathcal{T} \in \Omega} \int_{\mathbf{I}} \mathbf{P}(\mathcal{T}, \mathbf{f})\mathrm{d}\mathbf{f}} = \frac{\int_{\mathbf{I}} \sum_{\mathcal{T} \in \Omega} \mathbf{P}(\mathcal{T}, \mathbf{f}, \eta_S \leq t)\mathrm{d}\mathbf{f}}{\int_{\mathbf{I}} \sum_{\mathcal{T} \in \Omega} \mathbf{P}(\mathcal{T}, \mathbf{f})\mathrm{d}\mathbf{f}}. \quad (3)$$

Since we have not closed formula for the quantities involved in the ratio above, it has to be numerically computed, for instance by uniformly sampling the fossil ages in $\mathbf{I}$. Being given $N$ uniform samples of fossil age vectors $\mathbf{f}_1, \ldots, \mathbf{f}_N$, the estimated ratio is

$$\frac{\sum_{i=1}^{N} \sum_{\mathcal{T} \in \Omega} \mathbf{P}(\mathcal{T}, \mathbf{f}, \eta_S \leq t)}{\sum_{i=1}^{N} \sum_{\mathcal{T} \in \Omega} \mathbf{P}(\mathcal{T}, \mathbf{f}_i)}.$$

Unfortunately, because of its variance, the estimator above requires millions of samples to converge in our dataset. We tackle this issue in the same way as in Didier and Laurin (2020), that is by performing importance sampling (see, e.g., Robert and Casella, 2013). Briefly, an importance sampling procedure estimates the integral $\int_{\mathbf{I}} \sum_{\mathcal{T} \in \Omega} \mathbf{P}(\mathcal{T}, \mathbf{f}, \eta_n = t)\mathrm{d}\mathbf{f}$ by sampling the fossil ages not uniformly but following a density $\mathrm{q}^\star$ biased toward the ages corresponding to high values of $\sum_{\mathcal{T} \in \Omega} \mathbf{P}(\mathcal{T}, \mathbf{f}, \eta_n = t)$. We have basically that

$$\int_{\mathbf{I}} \sum_{\mathcal{T} \in \Omega} \mathbf{P}(\mathcal{T}, \mathbf{f}, \eta_S \leq t)\mathrm{d}\mathbf{f} = \int_{\mathbf{I}} \sum_{\mathcal{T} \in \Omega} \frac{\mathbf{P}(\mathcal{T}, \mathbf{f}, \eta_S \leq t)}{\mathrm{q}^\star(\mathbf{f})} \mathrm{q}^\star(\mathbf{f})\mathrm{d}\mathbf{f}.$$

It follows that if $\mathbf{f}'_1, \ldots, \mathbf{f}'_N$ are independently sampled from density $\mathrm{q}^\star$, this integral can be estimated by

$$\frac{1}{N} \sum_{i=1}^{N} \frac{\sum_{\mathcal{T} \in \Omega} \mathbf{P}(\mathcal{T}, \mathbf{f}'_i, \eta_S \leq t)}{\mathrm{q}^\star(\mathbf{f}'_i)}.$$

As in Didier and Laurin (2020), by noting that $\sum_{\mathcal{T} \in \Omega} \mathbf{P}(\mathcal{T}, \mathbf{f}', \eta_S \leq t)$ is bounded by (and strongly related to) $\sum_{\mathcal{T} \in \Omega} \mathbf{P}(\mathcal{T}, \mathbf{f}')$, we consider a density $\mathrm{q}^\star(\mathbf{f})$ proportional to $\sum_{\mathcal{T} \in \Omega} \mathbf{P}(\mathcal{T}, \mathbf{f})$ for our importance sampling procedure. Namely, we set

$$\mathrm{q}^\star(\mathbf{f}') = \frac{\sum_{\mathcal{T} \in \Omega} \mathbf{P}(\mathcal{T}, \mathbf{f}')}{\int_{\mathbf{I}} \sum_{\mathcal{T} \in \Omega} \mathbf{P}(\mathcal{T}, \mathbf{f})\mathrm{d}\mathbf{f}}.$$

Given $N$ iid samples $\mathbf{f}'_1, \ldots, \mathbf{f}'_N$ obtained with the density $\mathrm{q}^\star$ just above, the ratio of Equation 3 is estimated by

$$\begin{aligned}
\frac{\frac{1}{N} \sum_{i=1}^{N} \frac{\sum_{\mathcal{T} \in \Omega} \mathbf{P}(\mathcal{T}, \mathbf{f}'_i, \eta_S \leq t)}{\mathrm{q}^\star(\mathbf{f}'_i)}}{\int_{\mathbf{I}} \sum_{\mathcal{T} \in \Omega} \mathbf{P}(\mathcal{T}, \mathbf{f})\mathrm{d}\mathbf{f}} &= \frac{1}{N} \sum_{i=1}^{N} \frac{\sum_{\mathcal{T} \in \Omega} \mathbf{P}(\mathcal{T}, \mathbf{f}'_i, \eta_S \leq t)}{\sum_{\mathcal{T} \in \Omega} \mathbf{P}(\mathcal{T}, \mathbf{f}'_i)} \\
&= \frac{1}{N} \sum_{i=1}^{N} \frac{\sum_{\mathcal{T} \in \Omega} \mathbf{P}(\mathcal{T}, \mathbf{f}'_i) \prod_{n \in S} \frac{\mathbf{P}(0, t - \ell_{\mathbf{f}'_i, n})}{\mathbf{P}(0, T - \ell_{\mathbf{f}'_i, n})}}{\sum_{\mathcal{T} \in \Omega} \mathbf{P}(\mathcal{T}, \mathbf{f}'_i)} \\
&= \frac{1}{N} \sum_{i=1}^{N} \prod_{n \in S} \frac{\mathbf{P}(0, t - \ell_{\mathbf{f}'_i, n})}{\mathbf{P}(0, T - \ell_{\mathbf{f}'_i, n})}.
\end{aligned}$$

Since it is not possible to directly sample following the density $\mathrm{q}^\star$, which is known only up to the intractable normalization constant $\int_{\mathbf{I}} \sum_{\mathcal{T} \in \Omega} \mathbf{P}(\mathcal{T}, \mathbf{f})\mathrm{d}\mathbf{f}$, we proceed in the same way as Didier and Laurin (2020) by using the Metropolis-Hastings algorithm (e.g., Robert and Casella, 2013) to build a Markov chain with target density $\mathrm{q}^\star$. Let us recall the details of this procedure. Given the $i^{\text{th}}$ vector of fossil ages $\mathbf{f}'_i$ of this chain, we generate a candidate vector $\tilde{\mathbf{f}}$ first by uniformly selecting a fossil $f$ among all the fossils of the dataset and by replacing the corresponding entry of $\mathbf{f}'_i$ by a uniform sample in the interval centered on this entry of width equal to $\alpha$ time the with of the interval $\mathrm{I}_f$ that represents the uncertainty on the age of fossil $f$ (i.e., we use a usual sliding window proposal with reflection; $\alpha$ is a user-defined parameter). This proposal transition kernel is basically symmetrical, i.e., getting the candidate vector

$\tilde{\mathbf{f}}$ from $\mathbf{f}'_i$ has the same probability as getting the candidate vector $\mathbf{f}'_i$ from $\tilde{\mathbf{f}}$. The next fossil age vector $\mathbf{f}'_{i+1}$ in the Markov chain is

$$\mathbf{f}'_{i+1} = \begin{cases} \tilde{\mathbf{f}} & \text{with probability } \min\left\{\frac{\sum_{\mathcal{T}\in\Omega}\mathbf{P}(\mathcal{T},\tilde{\mathbf{f}})}{\sum_{\mathcal{T}\in\Omega}\mathbf{P}(\mathcal{T},\mathbf{f}'_i)},1\right\} \text{ and} \\ \mathbf{f}'_i & \text{with probability } 1-\min\left\{\frac{\sum_{\mathcal{T}\in\Omega}\mathbf{P}(\mathcal{T},\tilde{\mathbf{f}})}{\sum_{\mathcal{T}\in\Omega}\mathbf{P}(\mathcal{T},\mathbf{f}'_i)},1\right\}. \end{cases}$$

In the case where the speciation, extinction and fossilization rates of the model are not given *a priori* (e.g., estimated by maximum likelihood), we sum over all their possible values by assuming that they all follow the improper uniform distribution over $[0, +\infty]$ and by following a similar importance sampling procedure. In plain English, the importance sampling is now performed over both the fossil age vector $\mathbf{f}$ and the model rates $\Theta = (\lambda, \mu, \psi)$ by using the biased density $q^\star(\mathbf{f}, \Theta)$ proportional to $\sum_{\mathcal{T}\in\Omega}\mathbf{P}_\Theta(\mathcal{T}, \mathbf{f}')$, where $\mathbf{P}_\Theta(\mathcal{T}, \mathbf{f}')$ is the probability of the phylogeny with fossils $\mathcal{T}, \mathbf{f}'$ under the FBD model with rates $\Theta = (\lambda, \mu, \psi)$.

The proposal mechanism of the Markov chain used to sample according to this density first draws a Bernoulli random variable with a probability $\beta$ (provided by the user) according to which either a change in a fossil age or a change in the speciation, extinction or fossilization rate is proposed. The change in a fossil age is proposed in the exact same way as just above. The change in a rate is proposed first by selected randomly the speciation, extinction or fossilization rate according to user-defined probabilities then by using a sliding window of fixed length centered on the current selected rate.

The densities presented in the results section were obtained from the MCMC procedure presented just above by discarding the first 10000 iterations, and by keeping only 1 iteration over 200 for the subsequent iterations. We consider a window of width equal to 80% of the length of the age interval $\mathrm{I}_f$ to propose a change in the age of fossil $f$ and windows of size 0.5 to propose changes in the speciation, extinction and fossilization rates respectively. We propose a change in a fossil age with probability 0.75 and a change in the speciation, extinction or fossilization rate otherwise (the rate to be changed among the the speciation, extinction and fossilization rate is then drawn uniformly). We run the MCMC procedure until get 10000 iterations. The iterations thus obtained pass all the convergence tests of the `coda` R package with an effective size greater than 1000 for all the parameters (fossil ages and rates, Plummer *et al.*, 2006). All the parameters of the MCMC were manually tuned in order to get a satisfying behavior. They can be modified by calling the corresponding R function in the package.

### SI-4 Posterior probability density distributions of speciation, extinction and fossilization rates from the simulation study

Figure SI-1 displays the aggregated posterior densities of the speciation, extinction and fossilization rates corresponding to the five runs of simulations presented in the section "Simulation study". Let us recall that we performed five runs of 1000 simulations, all with speciation and extinction rates 0.2 and 0.19 and with fossilization rates 0.005, 0.01, 0.1, 1 and 5 respectively. For each of the five fossilization rates, we plotted the average posterior density of the speciation, extinction and fossilization rates over the 1000 simulated trees.

We observe that each density is roughly centered on the corresponding rate value used to simulate the data and that the distributions of the speciation and extinction rates become tighter as the fossilization rate increases.

### SI-5 Simulation study with two other levels of risk

We performed the exact same protocol as that described in Section "Simulation study" by considering confidence upper bounds at two other orders: 50% (Tables SI-1 and SI-2) and 75% (Tables SI-3 and SI-4). Tables SI-1 and SI-3 display the error percentage, that is the percentage of cases where the "real" extinction date is posterior to the upper bound provided by the six assessed methods. Tables SI-2 and SI-4 show the mean confidence interval widths obtained with each method at orders 50% and 75% respectively. For both orders 50% and 75%, we observe the same general behavior as that observed with the 95% confidence upper bounds.

| Fos. rate | **S&S** | **McI** | **Alr** | **Glo** | | **Int** | | | **Ref** | | |
|---|---|---|---|---|---|---|---|---|---|---|---|
| 0.005 | 29.29 | 36.29 | 11.34 | 31.24 | *32.73* | 44.91 | *48.15* | **49.59** | 47.44 | *49.92* | **50.12** |
| 0.01 | 34.68 | 42.23 | 13.97 | 32.72 | *34.67* | 49.39 | *50.43* | **51.16** | 50.10 | *50.17* | **50.06** |
| 0.1 | 44.15 | 51.75 | 11.85 | 43.50 | *43.19* | 51.31 | *51.36* | **51.62** | 50.38 | *50.15* | **50.15** |
| 1 | 49.97 | 54.38 | 6.32 | 50.00 | *50.02* | 50.65 | *50.68* | **50.80** | 50.49 | *50.46* | **50.46** |
| 5 | 49.79 | 51.87 | 2.09 | 49.59 | *49.65* | 49.67 | *49.74* | **49.78** | 49.59 | *49.66* | **49.66** |

Table SI-1: Error percentages obtained from the 50% confidence upper bounds provided by the six methods on the simulated extinct taxa where the **S&S**, **McI** and **Alr** computations were feasible (plain text), on those belonging to a simulated tree where a global fossil recovery rate can be estimated (italics; only for methods **Glo**, **Ref** and **Int**) and on all the extinct taxa (bold; only for methods **Ref** and **Int**).

| Fos. rate | **S&S** | **McI** | **Alr** | **Glo** | | **Int** | | | **Ref** | | |
|---|---|---|---|---|---|---|---|---|---|---|---|
| 0.005 | 13.3 | 9.4 | 36.0 | 9.3 | *9.5* | 4.2 | *4.1* | **3.9** | 3.7 | *3.7* | **3.7** |
| 0.01 | 10.0 | 7.1 | 28.9 | 7.1 | *7.1* | 3.6 | *3.5* | **3.4** | 3.4 | *3.4* | **3.4** |
| 0.1 | 3.2 | 2.4 | 14.0 | 2.4 | *2.4* | 1.9 | *1.8* | **1.8** | 1.9 | *1.9* | **1.9** |
| 1 | 0.6 | 0.5 | 4.7 | 0.5 | *0.5* | 0.5 | *0.5* | **0.5** | 0.5 | *0.5* | **0.5** |
| 5 | 0.1 | 0.1 | 1.6 | 0.1 | *0.1* | 0.1 | *0.1* | **0.1** | 0.1 | *0.1* | **0.1** |

Table SI-2: Mean confidence interval width in million years, i.e. the mean difference between the 50% confidence upper bounds provided by the three methods and the most recent fossil, on the simulated extinct taxa where the **S&S**, **McI** and **Alr** computations were feasible (plain text), on those belonging to a simulated tree where a global fossil recovery rate can be estimated (italics; only for methods **Glo**, **Int** and **Ref**) and on all the extinct taxa (bold; only for methods **Int** and **Ref**).

| Fos. rate | **S&S** | **McI** | **Alr** | **Glo** | | **Int** | | | **Ref** | | |
|---|---|---|---|---|---|---|---|---|---|---|---|
| 0.005 | 13.74 | 23.20 | 11.34 | 16.59 | *18.59* | 22.75 | *24.62* | **25.26** | 24.95 | *25.25* | **25.19** |
| 0.01 | 17.46 | 28.65 | 13.97 | 16.24 | *18.50* | 25.56 | *26.34* | **26.52** | 25.75 | *25.56* | **25.46** |
| 0.1 | 21.06 | 34.04 | 11.85 | 20.18 | *20.51* | 26.04 | *26.17* | **26.26** | 25.18 | *25.06* | **25.06** |
| 1 | 24.87 | 33.23 | 6.32 | 24.42 | *24.65* | 25.08 | *25.33* | **25.37** | 24.84 | *25.03* | **25.03** |
| 5 | 25.12 | 29.24 | 2.09 | 25.13 | *25.19* | 25.15 | *25.21* | **25.23** | 25.08 | *25.13* | **25.13** |

Table SI-3: Error percentages obtained from the 75% confidence upper bounds provided by the six methods on the simulated extinct taxa where the **S&S**, **McI** and **Alr** computations were feasible (plain text), on those belonging to a simulated tree where a global fossil recovery rate can be estimated (italics; only for methods **Glo**, **Ref** and **Int**) and on all the extinct taxa (bold; only for methods **Ref** and **Int**).

### SI-6 Divergence time and posterior probability density distributions from the empirical dataset

#### Probability density distributions of divergence times

We applied the approach presented in Didier and Laurin (2020) to the empirical dataset used here to compute the divergence time densities of the tree displayed in Figure 3 of the main document. Figure SI-2 displays this tree with the divergence time densities associated to all its internal nodes. The denser fossil record and our revisions of the ages of fossils based on recent literature yields slightly more recent divergence times (Fig. SI-2) than our previous study (Didier and Laurin, 2020, fig. 6).

Figure SI-3 presents the probability density of the divergence time between therapsids and sphenacontids. Didier and Laurin (2020) estimated that the divergence between sphenacodontids and therapsids

| Fos. rate | **S&S** | **McI** | **Alr** | **Glo** | | **Int** | | | **Ref** | | |
|---|---|---|---|---|---|---|---|---|---|---|---|
| 0.005 | 39.5 | 18.7 | 36.0 | 18.7 | *19.0* | 9.8 | *9.5* | **9.3** | 9.0 | *9.0* | **9.0** |
| 0.01 | 29.5 | 14.1 | 28.9 | 14.3 | *14.2* | 8.2 | *8.0* | **7.9** | 8.0 | *8.0* | **8.0** |
| 0.1 | 9.0 | 4.8 | 14.0 | 4.8 | *4.8* | 3.9 | *3.9* | **3.9** | 4.0 | *4.0* | **4.0** |
| 1 | 1.5 | 1.0 | 4.7 | 1.0 | *1.0* | 1.0 | *1.0* | **1.0** | 1.0 | *1.0* | **1.0** |
| 5 | 0.3 | 0.3 | 1.6 | 0.3 | *0.3* | 0.3 | *0.3* | **0.3** | 0.3 | *0.3* | **0.3** |

Table SI-4: Mean confidence interval width in million years, i.e. the mean difference between the 75% confidence upper bounds provided by the three methods and the most recent fossil, on the simulated extinct taxa where the **S&S**, **McI** and **Alr** computations were feasible (plain text), on those belonging to a simulated tree where a global fossil recovery rate can be estimated (italics; only for methods **Glo**, **Int** and **Ref**) and on all the extinct taxa (bold; only for methods **Int** and **Ref**).

dated from about 313 Ma (early Moscovian), but our current analysis places this divergence closer to 301 Ma, in the mid-Gzhelian (Fig. SI-3). Again, the more recent estimated age reflects the denser fossil record.

#### Posterior probability density distributions of rates and fossil ages

Figure SI-4 displays some posterior probability density distributions obtained from the MCMC sampling used to compute Figure SI-3. Since the fossil ages are given as time intervals in the empirical dataset, they have to be sampled as presented in Appendix SI-3. On top of the posterior distributions of the speciation, extinction and fossilization rates, we plotted those of a selection of fossil ages. Among these, we first observe that, though two fossils of *Dimetrodon limbatus* are associated to the time interval $(297, 290)$, the posterior densities of their age are negligible for ages greater than 293 Ma. Since the posterior density associated to an age value is proportional to the density of the phylogenetic tree when the corresponding fossil age is set to this value, this explains why the density of the speciation time of this taxon is negligible before 293 Ma, as shown in Figure SI-2. The posterior probability density distributions of fossils ages show various shapes. In particular, unlike that of *Dimetrodon limbatus*, some of them are almost uniform (e.g., that of *Tetraceratops insignis*).

Figure SI-4 shows that the posterior density of the fossilization rate supports rates about ten times higher than those observed with our dataset of Didier and Laurin (2020). This presumably reflects the incorporation of a fairly large amount of fossil occurrence data from the Paleobiology Database, which resulted in the incorporation of many additional occurrences of several taxa (but no additional taxa), as well as the more recent divergence times, which result in shorter branches. The slightly lower speciation and extinction rates observed in Figure SI-4 may simply reflect the fact that the taxonomic sample is not the same (the current study focuses on a subset of the clades included in our previous studies).

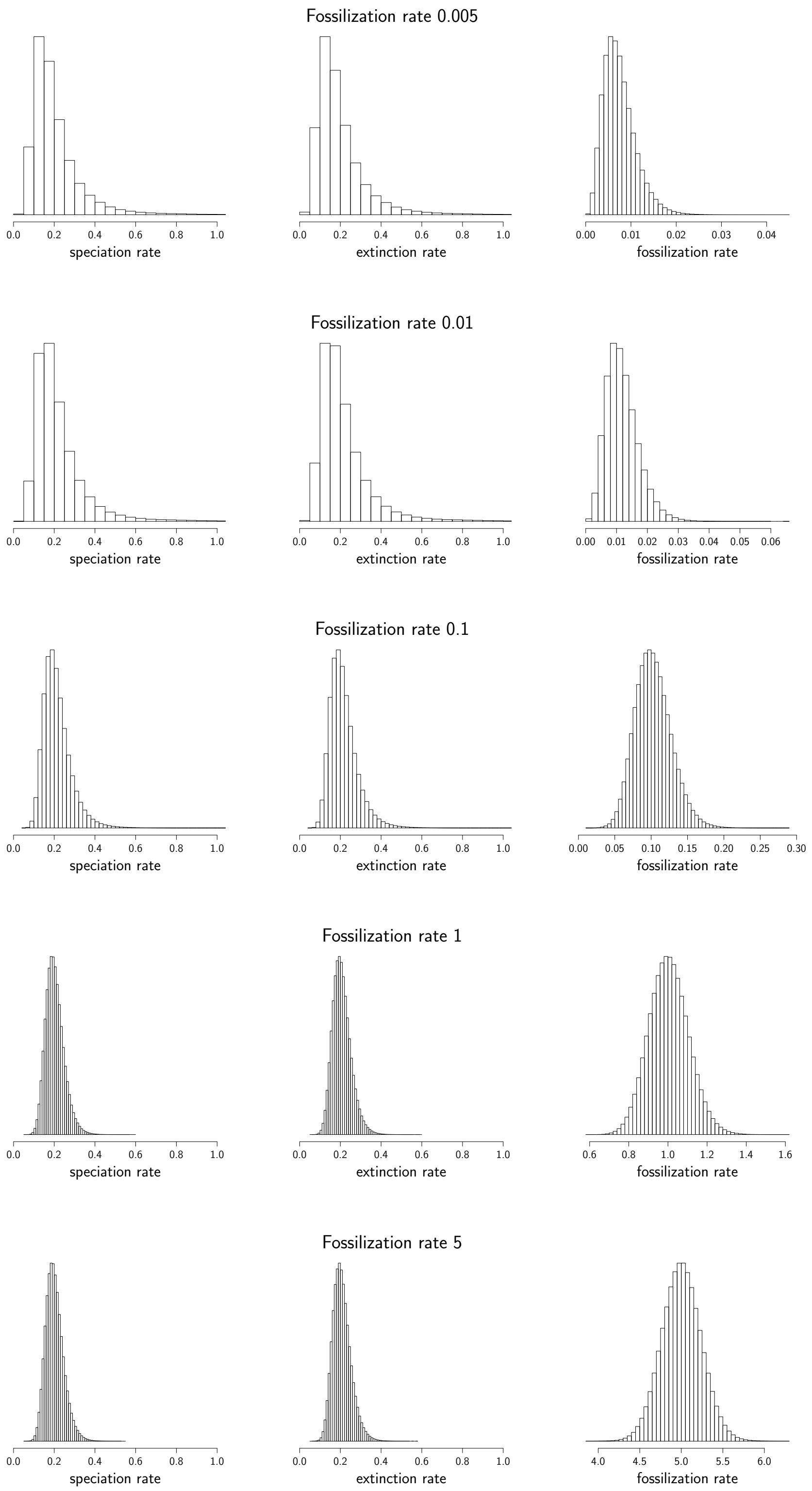


Figure SI-1: Posterior probability density distributions of the speciation, extinction and fossilization rates obtained from the MCMC procedure used to compute the upper bounds of extinction dates with the method **Int** (i.e., when the FBD rates are unknown) in our simulation study.

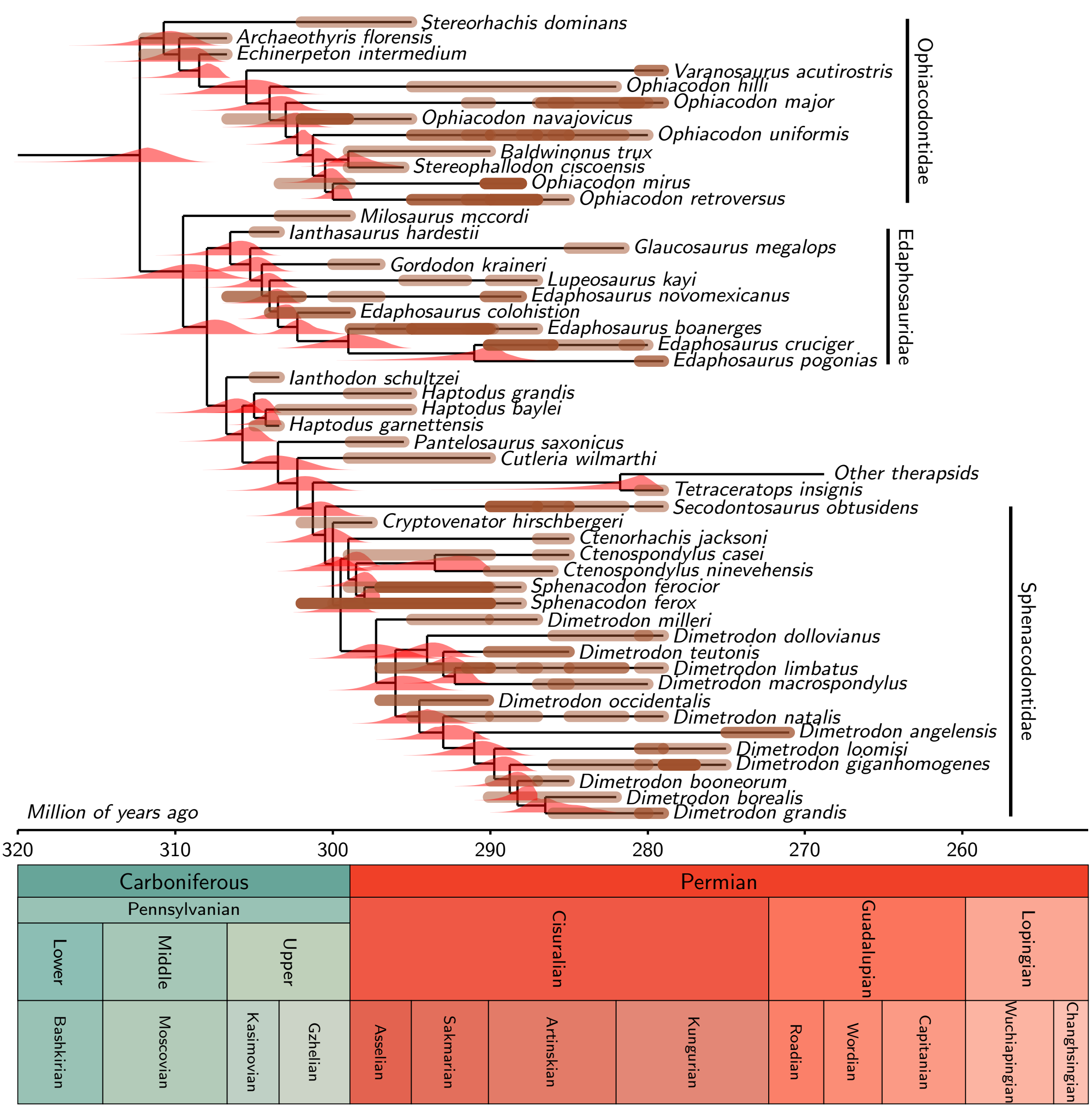


Figure SI-2: One of the 100 equally parsimonious trees of our dataset with the probability density distributions of its speciation times. Intervals of possible ages for each fossil are represented as thick brown segments with a certain level of transparency (darker brown segments correspond to overlapping intervals). The confidence interval on some fossil records extends below their basal node, most noticeably for *Dimetrodon limbatus*. With this tree topology, the oldest part of the range of possible ages of fossils of some taxa leads to very low probability densities of the whole tree with fossil; thus, it does not contribute significantly to the speciation time density distributions (Fig. SI-4). All nodes are older than the minimal age of all fossils, as they should be. The denser sample of the fossil record yields younger divergence times for eupelycosaur clades than those of our previous study (Didier and Laurin, 2020). This is because confidence intervals typically become narrower as sample size increases, and this phenomenon also applies to divergence times; as the fossil record becomes denser, our model adds less unobserved lineages below the first occurrence of a fossil on each lineage.

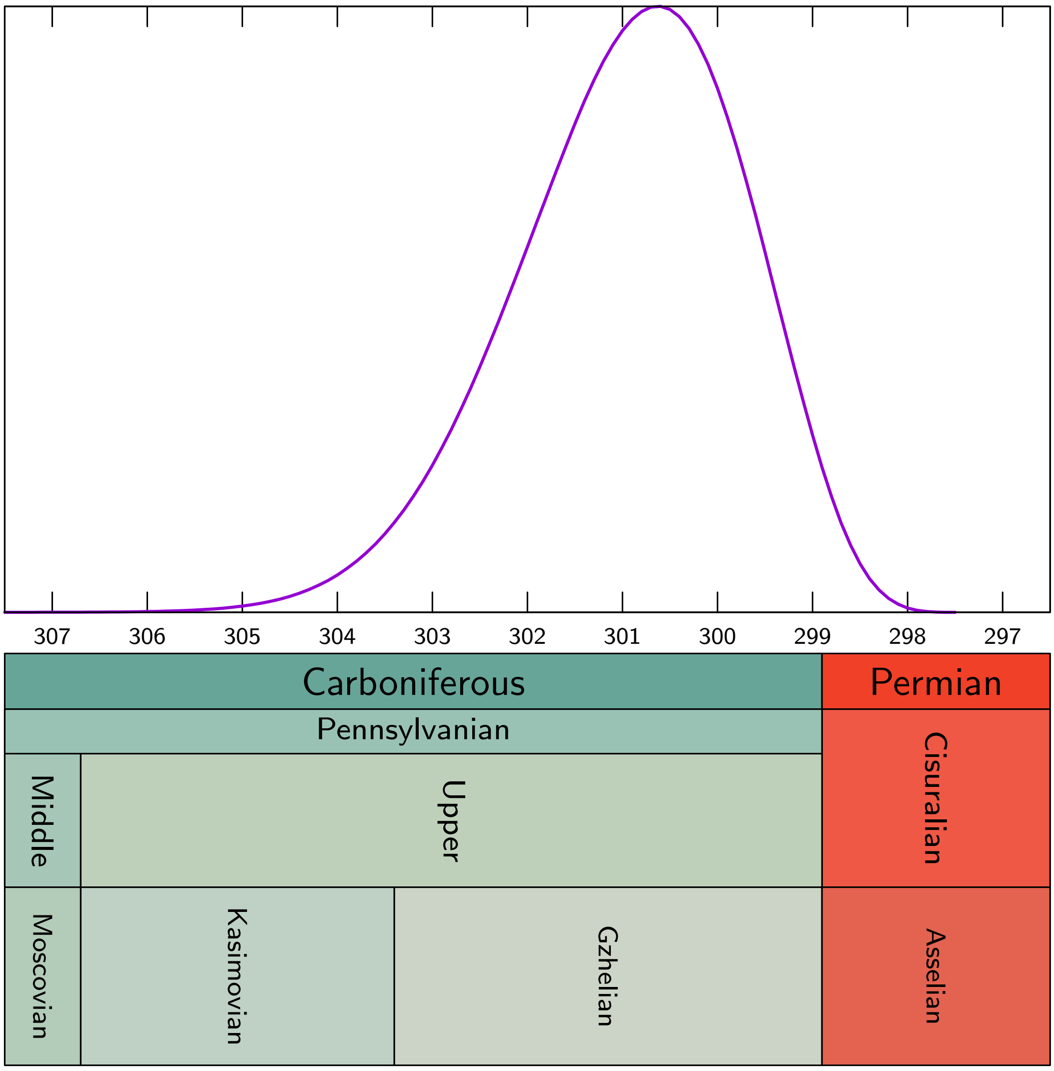


Figure SI-3: Probability density distribution of the age of the divergence between therapsids and sphenacontids.

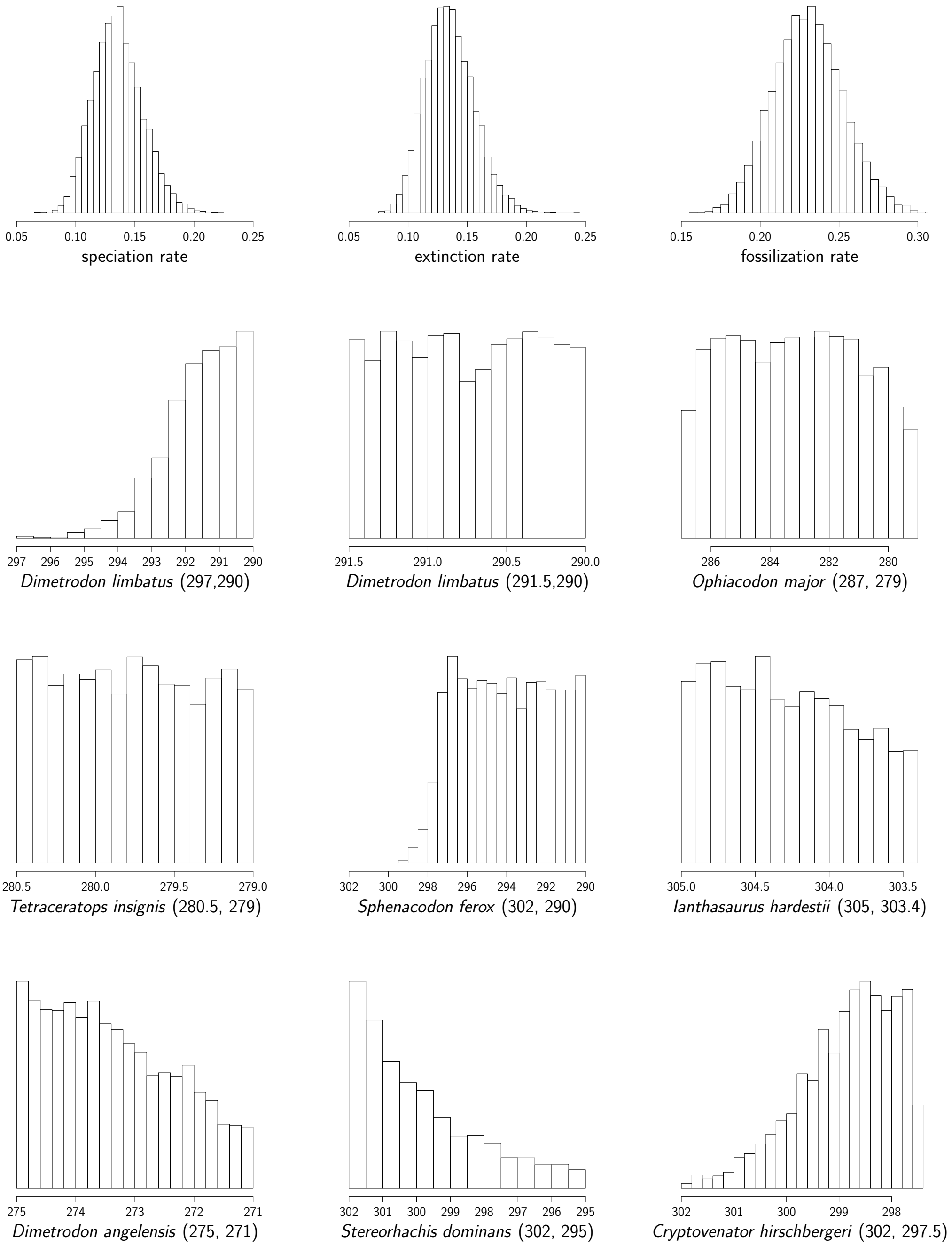


Figure SI-4: Posterior probability density distributions of the speciation, extinction and fossilization rates and of a selection of fossil ages obtained from the MCMC sampling used to compute the speciation distribution of Fig. SI-3.
